# Latitudinal variation in circadian rhythmicity in *Nasonia vitripennis*

**DOI:** 10.1101/594606

**Authors:** Silvia Paolucci, Elena Dalla Benetta, Lucia Salis, David Doležel, Louis van de Zande, Leo W. Beukeboom

**Affiliations:** Groningen Institute for Evolutionary Life Sciences, University of Groningen, P.O.Box 11103, 9700 CC, Groningen, The Netherlands; Institute of Entomology, Biology Center of the Czech Academy of Sciences, 370 05, Ceske Budejovice, Czech Republic

**Keywords:** circadian clock, latitudinal cline, local adaptation, free running period, locomotor activity

## Abstract

Many physiological processes of living organisms show circadian rhythms, governed by an endogenous clock. This clock has a genetic basis and is entrained by external cues such as light and temperature. Other physiological processes exhibit seasonal rhythms, that are also responsive to light and temperature. We previously reported a natural latitudinal cline of photoperiodic diapause induction in the parasitic wasp *Nasonia vitripennis* in Europe and a correlated haplotype frequency for the circadian clock gene *period* (*per*). To evaluate if this correlation is reflected in circadian behaviour, we investigated circadian locomotor activity of seven populations from the cline. We found that the proportion of rhythmic males is higher than females in constant darkness, and that mating decreased rhythmicity of both sexes. Only for virgin females, the free running period (*τ*) increased weakly with latitude. Wasps from the most southern locality had an overall shorter free running rhythm and earlier onset, peak and offset of activity during the 24 h period, than wasps from the northernmost locality. We evaluate this variation in rhythmicity as a function of *period* haplotype frequencies in the populations and discuss its functional significance in the context of local adaptation.

## INTRODUCTION

The daily rotation of the Earth around its axis causes oscillating photoperiods that have led to the evolution of a large variety of activity patterns of organisms. Many behavioural and physiological activities, like mating, feeding and resting, show distinct oscillating rhythms with a peak of activity at a specific moment during the light-dark (LD) cycle. These are driven by an endogenous clock that is reset daily (entrained) by the prevailing LD cycles and runs with an intrinsic period of approximately 24 hours in constant darkness (DD) (Pittendrigh et al., 1991). The length of this endogenous rhythm is called the *free running period* (*τ*).

Day length (photoperiod) oscillates also seasonally and is an important cue for season-dependent behaviours, like migration in birds, hibernation in mammals and diapause in insects (reviewed in Bradshaw and Holzapfel, 2007). Additionally, daily photoperiods depend on latitude, being almost constant near the equator and increasing in yearly variation towards higher latitudes. Hence, depending on latitude, organisms will experience different photoperiods over the year. Given the sensitivity of the circadian clock to light-dark fluctuations, it is conceivable that it also plays a role in seasonal rhythmicity (Bunning, 1960). There is accumulating evidence for a role of the circadian clock genes in photoperiodism in many species (Dalla Benetta et al., 2019; Koštál, 2011; Mukai and Goto, 2016; Saunders, 2010; Urbanová et al., 2016) and several studies have shown that seasonal responses differ geographically as a result of variation in photoperiodic conditions (Hut et al., 2013). Nevertheless, it is still unclear whether the observed natural variation in photoperiodic response is controlled by specific circadian clock properties as a whole, such as the pace and the phase of the endogenous clock or by pleiotropy of individual clock genes (Dalla Benetta et al., 2019; Hut and Beersma, 2011)

Larval diapause of the parasitoid *Nasonia vitripennis* is maternally induced following a certain number of days (switch point) at a given critical photoperiod (CPP) and shows a robust clinal photoperiodic response (Paolucci et al., 2013; Saunders, 2013). Apparently, a clock mechanism is responsible for the timing and counting of the light-dark cycles necessary for proper starting of the photoperiodic response (Saunders, 2013). Under long photoperiods, the switch point to start inducing diapause occurs late in life or not at all (Saunders, 1968). Interestingly, haplotype frequency distribution of the circadian clock gene *period (per)* follows the observed cline in photoperiodic diapause induction (Paolucci et al., 2016, 2013).

To investigate if the observed correlation of *per* haplotypes with seasonal response is reflected in natural variation in circadian activities, that are known to be regulated by the gene *period* (Dalla Benetta et al. 2019), we analysed seven populations collected along a European cline from Corsica to northern Finland. We first tested variation in the free running period *τ* and then analysed the timing and level of locomotor activity for the most southern and most northern populations. Our data indicate that activity timing and average free-running rhythm differ between southern and northern lines of *N. vitripennis*, suggesting latitude-dependent effect on circadian clock consistent with clinal *per* haplotypes.

## MATERIALS AND METHODS

### Experimental lines

To study variation in locomotor activity in *Nasonia vitripennis*, we used isofemale lines established from natural field collected populations (Paolucci et al., 2013). These lines originated from seven European sampling locations (OUL (Finland, Oulu): 65°3⍰40.16⍰N, 25°31⍰40.80⍰E; TUR (Finland, Turku): 61°15⍰40.53⍰N, 22°13⍰23.96⍰E; LAT (Latvia): 56°51⍰22.56⍰N, 25°12⍰1.38⍰E; HAM (Germany, Hamburg): 53°36⍰23.62⍰N, 10°10⍰17.74⍰E; SCH (Germany, Schlüchtern): 50°19⍰56.10⍰N, 9°30⍰47.00⍰E; SWI (Switzerland): 46°44⍰9.14⍰N, 7°6⍰57.34⍰E; COR (France, Corsica): 42°22⍰40.80⍰N, 8°44⍰ 52.80⍰E). Wasps were maintained on *Calliphora* spp. pupae as hosts in mass culture vials under LD 18:06, 20 ± 1°C, to minimise diapause induction. For establishing free running periods under constant darkness (DD) 17-25 isofemale lines from each location and 4-8 individuals from each isofemale line were used (797 females and 715 males). As some individuals died before all data were collected, only 1072 individuals could be used for locomotor activity analysis: 548 females (163 virgin and 385 mated) and 544 males (122 virgin and 402 mated).

### Locomotor activity recording

To quantify animal movement over time, individuals were collected one day after emergence (mated group) or collected as pupae and allowed to develop into adults at room temperature (virgin group) and kept either at LD 16:08 or LD 08:16 based on experimental group. Adults were then individually transferred, without anesthetization, to glass tubes (diameter 5mm x height 70mm) that were half filled with an agar gel containing 30% sugar. Trikinetics *Drosophila* activity monitors 2 (DAM2)(www.trikinetics.com) were used for activity registration with 32 wasps per monitor. All assays were performed in light-tight boxes in temperature-controlled environmental chambers with 20°C and 50% humidity. The light source in the box consisted of white light with a maximum light intensity of about 200 lum/sqf. The Trikinetics system monitors how many times per minute each individual wasp interrupts an infrared light beam that passes through the centre of the glass tube. To determine free-running period under constant darkness (DD), wasps were first entrained under LD16:8 for 4 days and then subjected to 10 days of DD. To compare daily activity profile under long and short photoperiods, adult wasps were recorded for 10 days in either LD 16:08 or LD 08:16 regime.

### Data analysis and statistics

The raw locomotor activity data were first visualized with the program ActogramJ (Schmid et al., 2011); available at http://actogramj.neurofly.de. Double-plot actograms obtained with this software were eye inspected and dead animals were omitted from further analysis. Under constant darkness (DD) it was possible to measure the period of activity (*τ*) with periodogram analysis available in ActogramJ, which incorporates chi-square test (Sokolove and Bushell, 1978). Linear mixed effect models were used to test the effect of location, latitude, sex and mating status on the percentage of rhythmic individuals and on the length of the free running period *tau*, with isofemale line nested into location as random effect (package *nlme*). Only rhythmic individuals were included in these tests. All statistical analyses were performed with R statistical software (version 3.4.1, R Development Core Team 2012).

Under LD conditions the average activity was calculated as described by (Schlichting and Helfrich-Förster, 2015). The first 4 days of entrainment were excluded from the analysis. To determine the onset and offset of activity on each day data per wasp have to be plotted as bar diagrams with each bar representing the sum of activity within 20 min. The onset represents the first time when activity starts to rise consecutively, whereas the offset is when activity reaches the level, which is stable during the night phase. To determine the timing of the peaks, the data are smoothed by a moving average of 30 min. Through this process, randomly occurring spikes are reduced and the real maximum of the activity can be determined. The average phase of the onset, peak and offset, represented in *Zeitgeber time* (ZT, where ZT 0 represents the time when the light turned on), was compared between different lines and treatments.

## RESULTS

### Rhythmicity and free running periods (τ)

The percentage of rhythmic individuals ranged from 75-84% and did not show a latitudinal cline. Overall, there was a significant difference in the proportion of rhythmic males (0.90) and rhythmic females (0.68) (GLM, effect of location x sex: *χ*^*2*^ = 65.71, *P* < 0.01), with small but significant differences between locations (GLM, effect of location x sex: *χ*^*2*^ = 33.00, *P* < 0.01). Within each sex, there was a significant effect of location (GLM, effect of location: for females *χ*^*2*^= 13.28, *P* < 0.05; for males *χ*^*2*^ = 13.46, *P* < 0.05), but no significant correlation with latitude, both for the overall data (GLM, effect of latitude: *χ*^*2*^ = 1.77, *P* = 0.18) and for the sexes separately (GLM, effect of latitude: for females *χ*^*2*^ = 1.65, *P* = 0.19; for males *χ*^*2*^ = 0.16, *P* = 0.68).

There was a significant difference in rhythmicity between mated and virgin individuals. 58% of mated females were rhythmic compared to 92% of virgin females (GLM model, effect of mating status within females: *χ*^*2*^= 50.25, *P* < 0.01) (Fig. 1). For males, 97% of virgin individuals were rhythmic and 88% of mated ones (GLM model, effect of mating status within males: *χ*^2^ = 12.18, P < 0.01). There was, however, no interaction effect between mating status and sex (GLM, effect of mating status x sex: *χ*^*2*^ = 1.57, *P* = 0.20). Again, no latitudinal cline of rhythmicity was detected, when mating status was taken into account (GLM, effect of latitude: for virgin individuals *χ*^*2*^ = 2.22, *P* = 0.13; for mated individuals *χ*^*2*^ = 1.60, *P* = 0.20) (Fig. 2.). These results indicate that there is variation between populations, sexes and mating status in proportions of rhythmic individuals, but this variation does not follow a geographical cline.

**Fig. 1.**
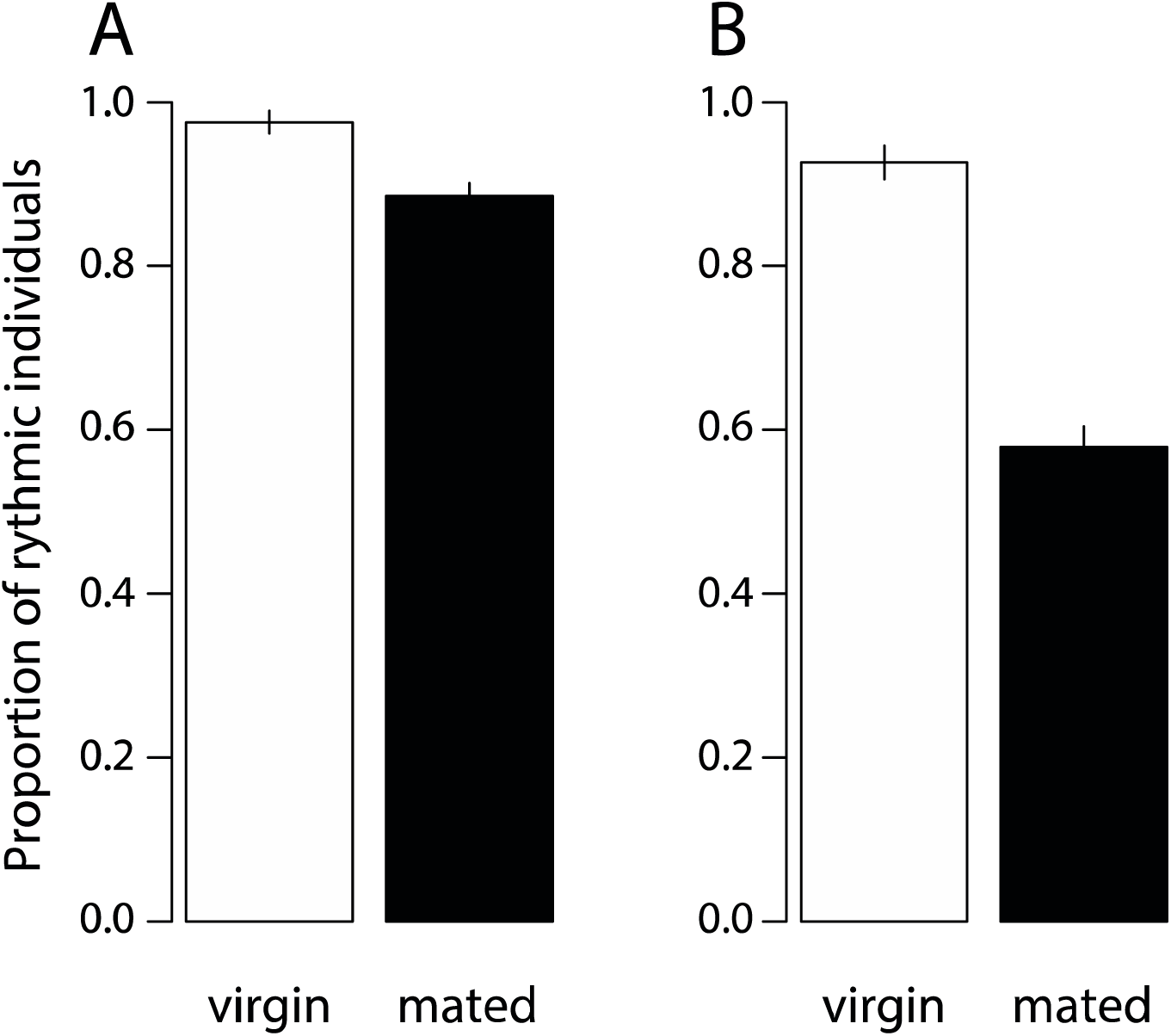
Effect of mating status on proportion of individuals with internal rhythmicity. Rhythmicity in *Nasonia vitripennis* individuals measured as proportion of individuals with internal rhythmicity under constant darkness in **(A)** males and **(B)** females. Rhythmic individuals are those individuals for whom a significant periodicity could be detected across several consecutive days in constant darkness and tau could be measured. Bars indicate standard errors. There is a significant difference between sexes and mating status (see text).

**Fig. 2.**
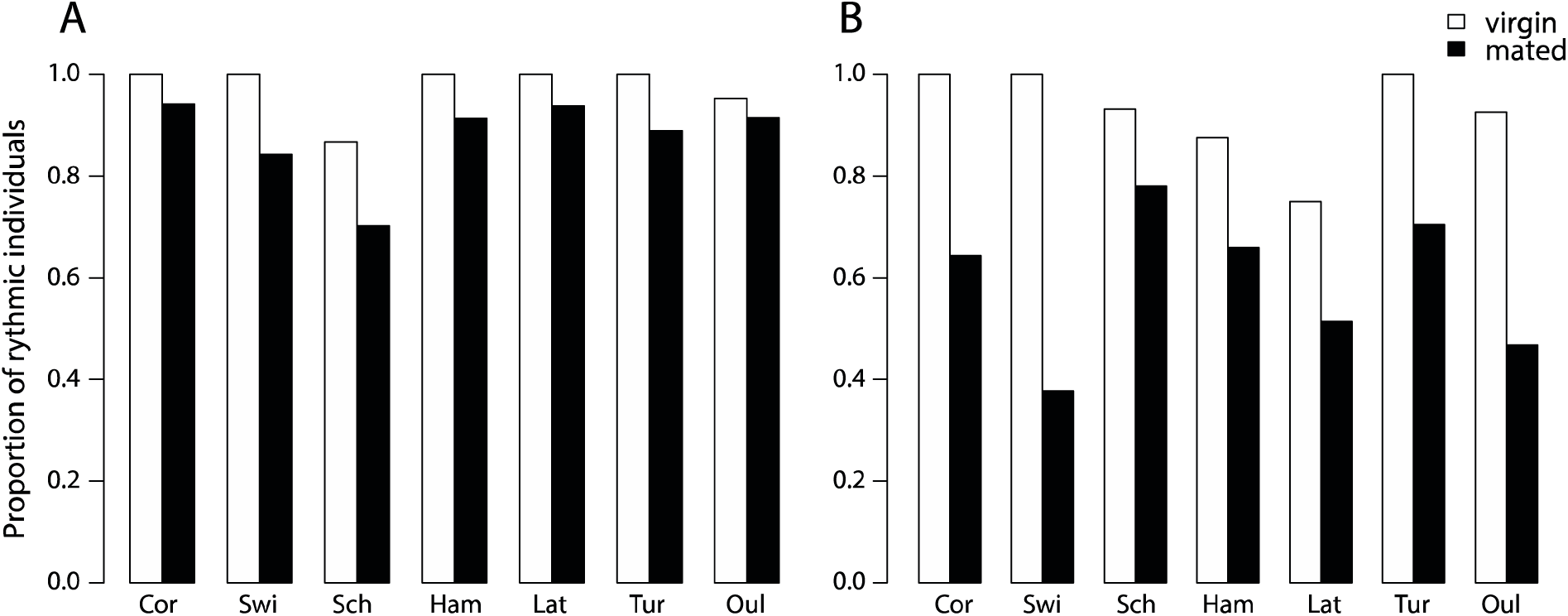
Rhythmicity under constant darkness of wasps originating from the European latitudinal transect. Proportion of rhythmic *Nasonia vitripennis* individuals in populations originating from seven locations in Europe in **(A)** males and **(B)** females. Locations along the x-axis are arranged from lower to higher latitude, see text for locality details.

Among all the tested individuals there was a large variation in *τ*, which ranged from 22 hours of the southern Corsica lines to 27.5 hours of the northern Oulu lines. Overall the free running period was shorter for individuals of southern latitudes and increased towards the north (Fig. 3). Females had a longer τ compared to males (24.39 ± 0.04 and 23.97 ± 0.04 h, respectively) and virgin individuals had longer τ compared to mated individuals (24.49 ± 0.05 and 24.00 ± 0.03 h, respectively). Sex, mating status and locality had significant effects on τ (LME model, effect of sex: *LRT* = 2.22, *P* = 0.13; effect of mating status: *LRT* = 44.32, *P* < 0.01; effect of location: *LRT* = 38.38, *P* < 0.01). The effect of location was significant for virgin females (LME, effect of location: *LRT* = 14.56, *P* < 0.05) and mated males (LME, effect of location: *LRT* = 13.02, *P* < 0.05) but not for mated females (LME effect of location: *LRT* = 12.02, *P* = 0.897) and virgin males (LME, effect of location: *LRT* = 4.19, *P* = 0.895). A very shallow but significant latitudinal cline in *τ* for virgin females (LME, effect of latitude: *LRT* = 4.28, r^2^ = 0.20, *P* < 0.001) was detected, but not for males and mated females (Fig. 3). The average *τ* for virgin COR females was 24.6 ± 0.12 h and for OUL 25.42 ± 0.11 h corresponding to a difference of 49.2 minutes.

**Fig. 3.**
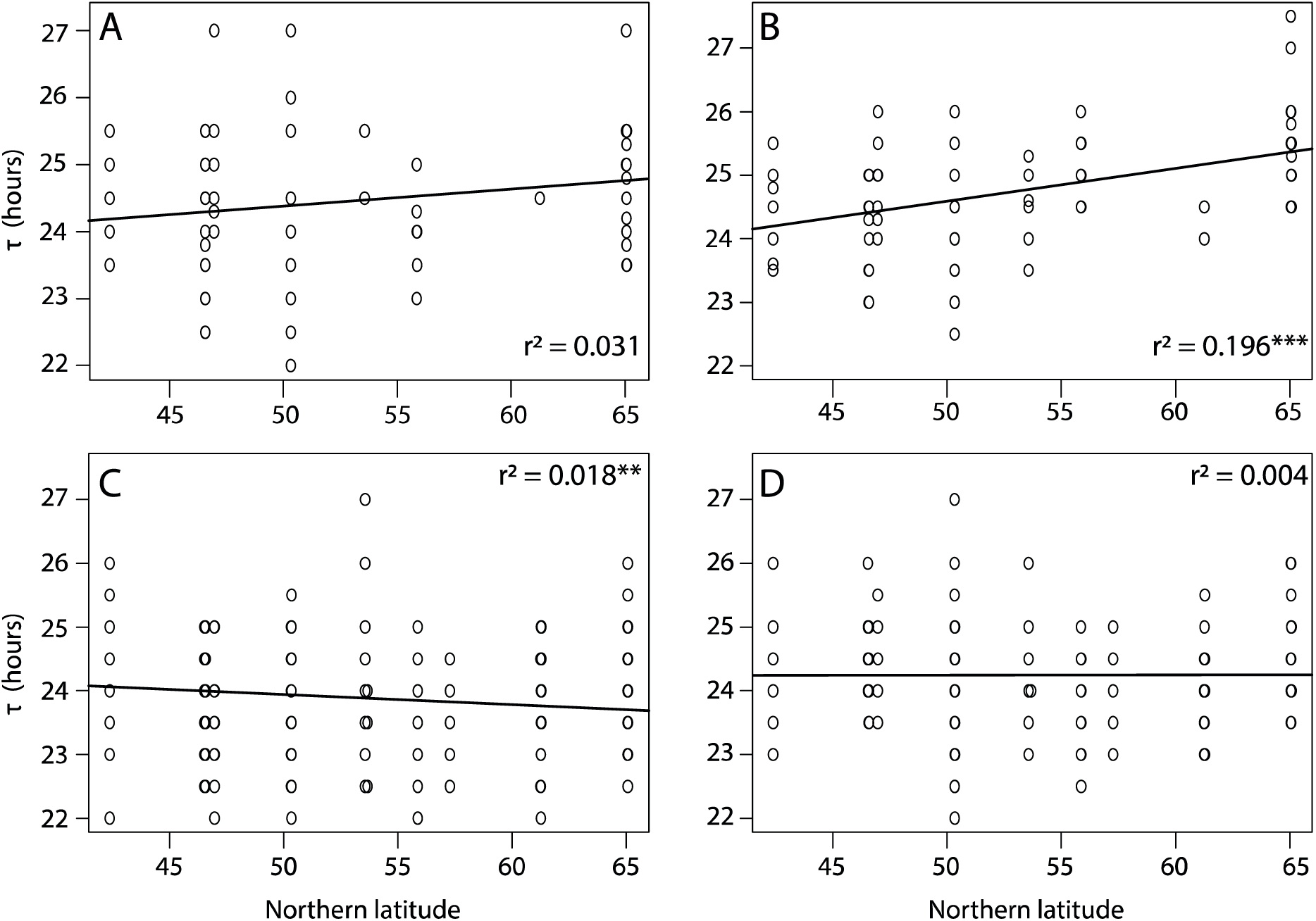
Free running period (τ) under constant darkness (DD) as a function of latitude. Free running period (τ) of **(A)** virgin males, **(B)** virgin females, **(C)** mated males and **(D)** mated females in *Nasonia vitripennis* populations collected along a latitudinal gradient in Europe. Asterisks indicate a significant effect of location along the cline (*** *P* < 0.001 and ** *P* < 0.05, linear mixed effect model)

### Activity timing

Virgin females from 5 isogenic lines established from populations of the two extremes of the sampling range were exposed to a light-dark regime of either LD16:08 or LD08:16h for 4 days. Under LD16:08, both southern and northern wasps displayed a unimodal activity pattern (Fig. 4), but with significant differences in the timing of onset, peak and offset of activity (Table 1, Table S1). Southern wasps started activity on average around ZT 0, which is about two hours earlier than northern wasps (Fig. 4, Table 1, Table S1). Southern wasps displayed maximum activity around ZT 5, while northern wasps peaked at ZT 8 (Fig. 4, Table 1, Table S1). Offset of activity was around ZT 13 and ZT 16 for southern and northern wasps, respectively (Fig. 4, Table 1, Table S1). Thus, southern wasps were more active in the first half of the light period and northern wasps towards the end of the day.

**Table 1:**
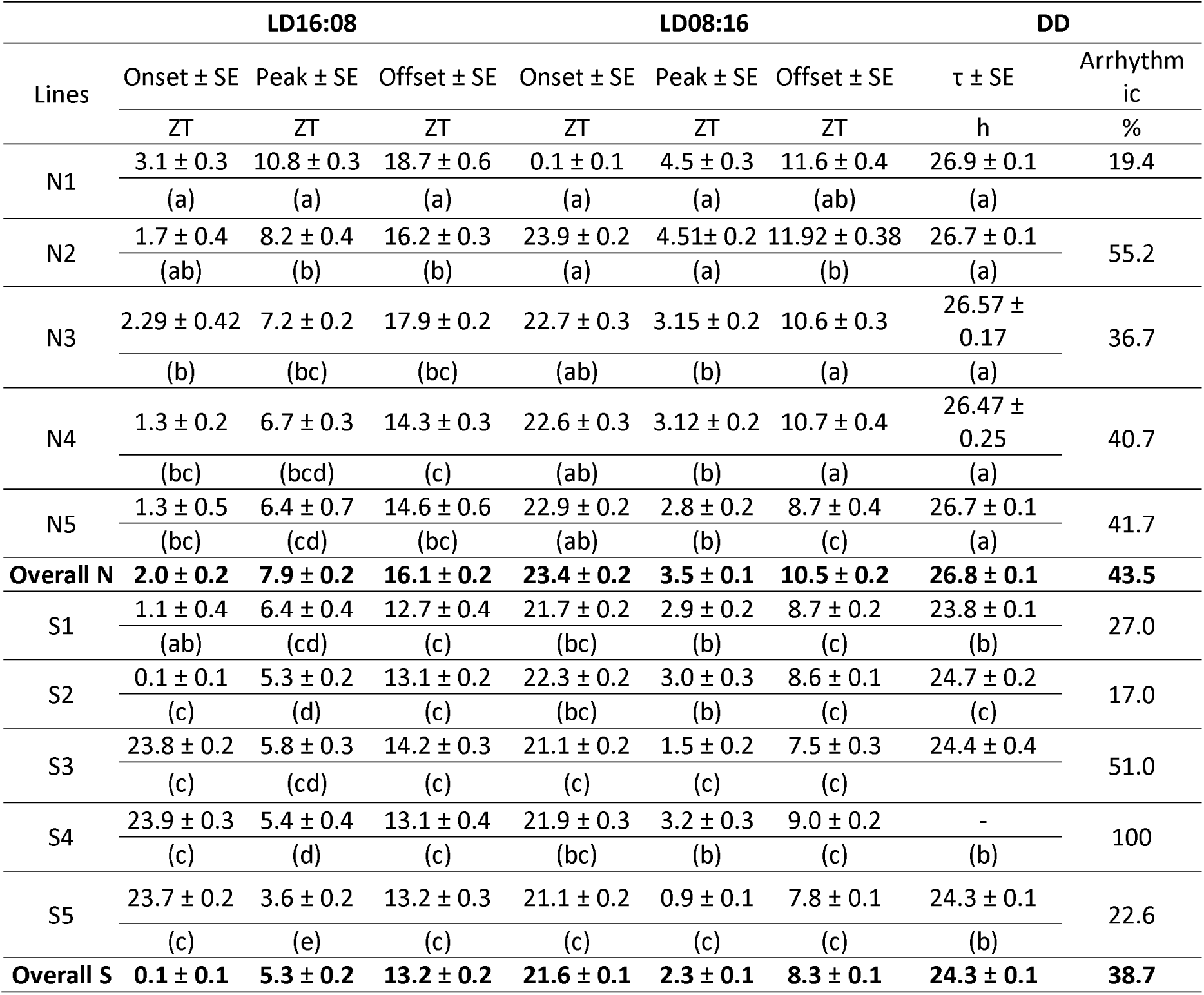
Timing of onset, peak and offset of activity for Corsica (45°) (S1, S2, S3, S4, S5) and Oulu (65°) (N1, N2, N3, N4, N5) wasps under long (LD16:08) and short (LD08:16) day conditions and free running rhythm (τ) under constant condition (DD). ZT (h) is zeitgeber time in hours. Different letters indicate significant differences (*P* < 0.05, ANOVA with a Tukey’s post hoc multiple-comparisons test).

**Fig. 4.**
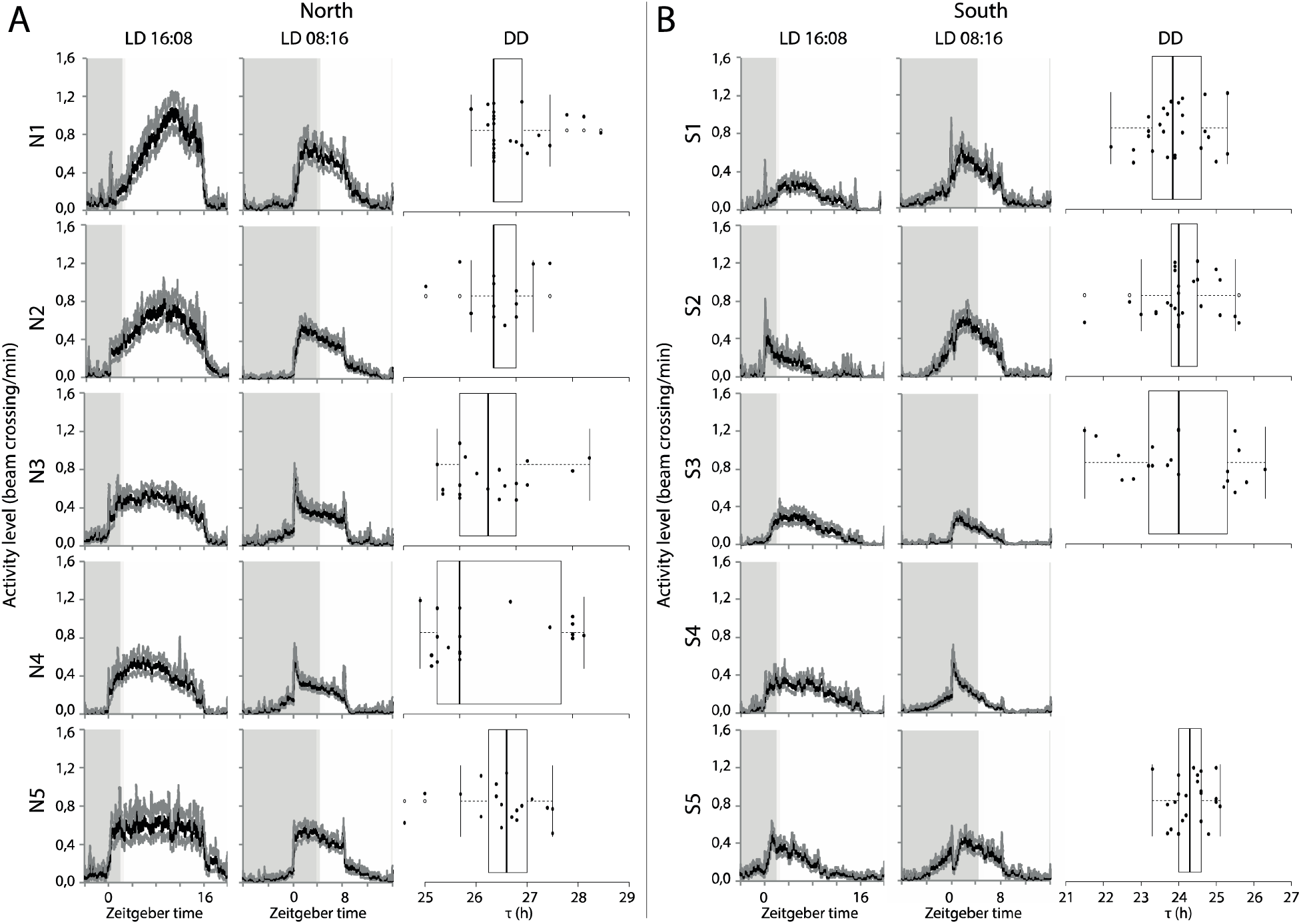
Locomotor activity of northern and southern wasps under different LD cycles. **(A)** Locomotor activity profiles for isogenic lines derived from Oulu (65°N) (N1, N2, N3, N4, N5) and **(B)** from Corsica (45°N) (S1, S2, S3, S4, S5) at long (LD16:08) and short (LD 08:16) day regimes. Grey shading indicates the night phase and white shading indicates the day phase. *Zeitgeber time* is indicated along the X-axis and ZT0 represents the time of light turn-on. Activity is calculated as average of bin crosses/minute of 25-32 individuals each over 24 h periods. Box plots represent the free running period (τ) in constant darkness (DD). Box plots depict the median (thick horizontal line within the box), the 25 and 75 percentiles (box margins) and the 1.5 interquartile range (thin horizontal line). Note that Line S4 is not rhythmic under DD. Note different scale on x-axis for A and B panels.

Under shorter photoperiod LD08:16 southern wasps started their activity also earlier than northern wasps (Fig. 4, Table 1, Table S2). Onset of activity occurred when the light was still off, around ZT 21.5. Northern wasps became active at ZT 0 (Fig. 4, Table 1, Table S2). The peaks of activity differed of about one-and-half hour, at ZT 2.5 and ZT 4 for southern and northern wasps, respectively (Fig. 4, Table 1, Table S2). Offset of activity was at ZT 8 for southern wasps. Northern wasps prolonged activity for more than two hours into darkness, until ZT 10.5 on average (Fig. 4, Table 1, Table S2). Thus, under both light regimes there is a difference in phase of activity between southern and northern wasps. The consequence is that in short photoperiod, southern wasps start activity in the dark and finish in the light phase, whereas northern wasps start activity at the beginning of the light phase and continue in the dark. Additionally, we observed a positive correlation (R2 = 0.51; *P* < 2e-16, linear mixed effect model) between the activity phase and free running rhythm of the wasps (Fig. S1).

In agreement with the results obtained for the isofemale lines, the southern and northern isogenic lines differed in τ under constant conditions (Fig. 4, Table1). The average free-running period of southern wasps was 24.3 ± 0.1h, which differed significantly from the longer τ of 26.7 ± 0.1h of the northern ones (*P* < 0.001).

## DISCUSSION

*Nasonia vitripennis* has a broad distribution and it is thus expected to exhibit natural variation in biological rhythms (Hut et al., 2013). Latitudinal cline variation in diapause induction (seasonal response), correlating with the clock gene *per*, is already reported by (Paolucci et al., 2016, 2013). Here we describe natural variation for several properties of circadian locomotor activity of *N. vitripennis*. Despite the proportion of rhythmic individuals being generally high, we observed significant differences between females and males: males are on average more rhythmic than females and have shorter free-running periods (*τ*). Similar differences between both sexes were observed by Floessner et al., (2019) and in the laboratory strain *N. vitripennis* AsymC (Bertossa et al., 2013). Virgin individuals of both sexes are highly rhythmic in constant darkness. The difference in rhythmicity between males and females is more apparent in mated individuals. Moreover, mated males maintain a high level of rhythmicity (although lower compared to virgin males), whereas most females lose their internal rhythmicity after mating. Similar effects of mating status on rhythmicity were found in the ant species *Camponotus compressus*, in which ovipositing queens exhibited arrhythmic locomotor activity during the egg laying phase and restored rhythmicity afterwards (Sharma et al., 2004). In addition, we found a significant interaction between locality and mating status on proportion of rhythmic individuals; for example, the effect of mating status in females of the SWI population is larger than in the SCH population. This might reflect standing genetic variation for rhythmic behaviour within and among populations. It is also possible that environmental factors affect the rhythmic locomotor activity as was recently shown for the northern fly species *Drosophila montana*, in which the proportion of rhythmic individuals was higher at lower temperature (Kauranen et al., 2012). It would be worthwhile to investigate how environmental conditions (temperature and light intensity among others) influence circadian locomotor activity behaviour of males and females in more insect species and test whether the level of variability will differ between populations from different geographical regions. Such studies could reveal differential selection pressures for stability of the circadian clock under different conditions.

We also found differences in the free running period (*τ*) between sexes and locations. Towards the south, wasps have a faster clock with *τ* close to 24h, whereas wasps from northern latitudes have a slower clock with *τ* longer than 24h. For virgin females, we observed a weak, but significant, latitudinal cline for τ, increasing towards higher latitudes. The presence of a positive latitudinal cline from south to north in DD rhythm was previously reported for *Drosophila* species (Prabhakaran and Sheeba, 2012, 2013), but only few studies have addressed variation of free running rhythms within a species. For example, in the model plant *Arabidopsis thaliana* the free running period under DD increases towards northern latitude, and correlates with clinal variation in seasonal flowering time regulated by photoperiodic cycles (Michael, 2003). In insects, similar results (*i*.*e*. longer *τ* towards northern latitude) were reported from the mosquito *Culex pipiens* (Shinkawa et al., 1994) and the linden bug *Pyrrhocoris apterus* (Pivarciova et al., 2016). This suggests that the latitudinal differences in free running period are the result of selection on traits that enable local adaptation. One possibility is selection for phase of activity in which a faster clock corresponds to earlier activity phase and a slower clock to later activity phase.

The period of the circadian clock of *Nasonia* females can reflect the timing of locomotor activity under LD conditions. Indeed, we observed a positive correlation between the activity phase and free running rhythm of the wasps (i.e. wasps with shorter *τ* have earlier activity pea; Fig. S1). Southern and northern wasps displayed profound differences in their daily locomotor activity. Southern wasps were mainly active in the morning with an increase of activity before the light turned on during short photoperiod, whereas northern ones present a unimodal evening activity, with a prolonged evening peak at the shorter photoperiod. This shifted activity pattern between southern and northern wasps can reflect local adaptation. In the south, temperatures are known to become high in the middle, late afternoon and shifting the activity to the coolest part of the day (the morning) might be a response of insects that live in a hot environment (Prabhakaran and Sheeba, 2013, 2012). In contrast, species that live at higher latitudes have to cope with lower temperatures and longer photoperiods (Helfrich-Förster et al., 2018). Indeed, the northern *Nasonia* lines have a reduced morning activity with their activity peak in the second part of the day when temperatures are higher. Similar differences in activity patterns between southern and northern populations have been reported for *Drosophila*, albeit between *Drosophila* species rather than populations within species (Prabhakaran and Sheeba, 2012, 2013; Helfrich-Förster et al., 2018). However, the overall activity profile of *N. vitripennis* is rather broad compared to the more precisely timed behaviour of *Drosophila melanogaster*, possibly reflecting a stronger selection on activity phase in *D. melanogaster* than *N. vitripennis*. On the other hand, *Nasonia* exhibits a stronger photoperiodic response (Saunders 1968). Moreover, the observed, albeit weak, cline in *τ* for virgin females, follows that of the frequencies of *period* haplotypes correlating with the cline for photoperiodic diapause induction (Paolucci et al., (2013, 2016) and the critical photoperiod to induce of *Nasonia* diapause was affected when *per* expression and *τ* were altered (Dalla Benetta et al., 2019; Floessner et al., 2019). If *per* participates in photoperiod measurement by fine-tuning critical day length to latitude-dependent requirements, this would suggest an involvement of this clock gene in the photoperiodic timer of *Nasonia*. Consequently, the cline in free running period would reflect a mere “side effect” of the selection pressure on seasonal rhythms. In agreement with this, recent work by Floessner et al. (2019) found a strong light resetting of the *Nasonia* circadian clock that allows wasps to entrain to a wide range of light-dark cycles, including the northern more extreme photoperiods, without negative effect on fitness.

In conclusion, we described natural variation in the period and phase of daily rhythms between southern and northern *N. vitripennis* lines. Many traits related to circadian activity show a high level of plasticity, which allows flexibility in daily activities depending on internal conditions (e.g. mating status) or external environmental conditions (e.g. light dark cycle, presence of food). Nevertheless, variation between geographic locations is maintained even in the plastic response to different stimuli, suggesting that natural selection acts on the response of the circadian system to the environment and not on the circadian clock *per se*. Clearly, more detailed functional experiments are required to reveal the exact molecular mechanism underpinning circadian clock, photoperiodic timer and their mutual connections.

## Acknowledgements

This work was funded by EU Marie Curie Initial Training Network INsecTIME (Grant Nr. 316790). We would like to thank all the participants of the network for helpful and stimulating discussion and the members of the Evolutionary Genetics, Development & Behaviour Group for discussion and advices on statistical analysis.

