## Supplementary information for "Latitudinal variation in circadian rhythmicity in *Nasonia vitripennis*"

**Table S1**: Statistical analysis of circadian timing between southern and northern *Nasonia vitripennis* under LD16:08. Indicated are *P*-values from ANOVA with a Tukey’s multiple-comparisons test. In bold *P* < 0.05.

|  | **Onset LD16:08** | | | | | | | | | | | | | | | | | |
| --- | --- | --- | --- | --- | --- | --- | --- | --- | --- | --- | --- | --- | --- | --- | --- | --- | --- | --- |
|  | N1 | N2 | N3 | | N4 | | N5 | | S1 | | S2 | | S3 | | S4 | | S5 | |
| N1 |  | 0.281 | 0.456 | | **< 0.001** | | **0.006** | | **< 0.001** | | **< 0.001** | | **< 0.001** | | **< 0.001** | | **< 0.001** | |
| N2 |  |  | 0.948 | | 0.995 | | 0.999 | | 0.938 | | **0.009** | | **< 0.001** | | **0.001** | | **< 0.001** | |
| N3 |  |  |  | | 0.364 | | 0.601 | | 0.139 | | **< 0.001** | | **< 0.001** | | **< 0.001** | | **< 0.001** | |
| N4 |  |  |  | |  | | 1.000 | | 0.999 | | 0.161 | | 0.239 | | **0.040** | | **0.010** | |
| N5 |  |  |  | |  | |  | | 0.999 | | 0.277 | | 0.060 | | 0.093 | | **0.031** | |
| S1 |  |  |  | |  | |  | |  | | 0.337 | | 0.065 | | 0.104 | | **0.030** | |
| S2 |  |  |  | |  | |  | |  | |  | | 0.997 | | 0.999 | | 0.988 | |
| S3 |  |  |  | |  | |  | |  | |  | |  | | 1.000 | | 1.000 | |
| S4 |  |  |  | |  | |  | |  | |  | |  | |  | | 0.999 | |
| S5 |  |  |  | |  | |  | |  | |  | |  | |  | |  | |
|  | **Peak LD16:08** | | | | | | | | | | | | | | | | | |
|  | N1 | N2 | | N3 | | N4 | | N5 | | S1 | | S2 | | S3 | | S4 | | S5 |
| N1 |  | **< 0.001** | | **< 0.001** | | **< 0.001** | | **< 0.001** | | **< 0.001** | | **< 0.001** | | **< 0.001** | | **< 0.001** | | **< 0.001** |
| N2 |  |  | | 0.4334 | | 0.05 | | **0.01** | | **0.008** | | **0.009** | | **< 0.001** | | **0.001** | | **< 0.001** |
| N3 |  |  | |  | | 0.989 | | 0.837 | | 0.878 | | **0.004** | | 0.138 | | **0.007** | | **< 0.001** |
| N4 |  |  | |  | |  | | 0.999 | | 0.999 | | 0.157 | | 0.791 | | 0.216 | | **< 0.001** |
| N5 |  |  | |  | |  | |  | | 1.000 | | 0.569 | | 0.988 | | 0.653 | | **< 0.001** |
| S1 |  |  | |  | |  | |  | |  | | 0.346 | | 0.950 | | 0.435 | | **< 0.001** |
| S2 |  |  | |  | |  | |  | |  | |  | | 0.994 | | 1.000 | | **0.009** |
| S3 |  |  | |  | |  | |  | |  | |  | |  | | 0.997 | | **< 0.001** |
| S4 |  |  | |  | |  | |  | |  | |  | |  | |  | | **0.010** |
| S5 |  |  | |  | |  | |  | |  | |  | |  | |  | |  |
|  | **Offset LD16:08** | | | | | | | | | | | | | | | | | |
|  | N1 | N2 | | N3 | | N4 | | N5 | | S1 | | S2 | | S3 | | S4 | | S5 |
| N1 |  | **< 0.001** | | **< 0.001** | | **< 0.001** | | **< 0.001** | | **< 0.001** | | **< 0.001** | | **< 0.001** | | **< 0.001** | | **< 0.001** |
| N2 |  |  | | 0.999 | | **0.031** | | 0.124 | | **< 0.001** | | **< 0.001** | | **0.011** | | **< 0.001** | | **< 0.001** |
| N3 |  |  | |  | | 0.127 | | 0.366 | | **< 0.001** | | **< 0.001** | | 0.055 | | **< 0.001** | | **< 0.001** |
| N4 |  |  | |  | |  | | 0.999 | | 0.053 | | 0.272 | | 0.999 | | 0.337 | | 0.463 |
| N5 |  |  | |  | |  | |  | | 0.021 | | 0.132 | | 0.999 | | 0.173 | | 0.256 |
| S1 |  |  | |  | |  | |  | |  | | 0.998 | | 0.143 | | 0.998 | | 0.986 |
| S2 |  |  | |  | |  | |  | |  | |  | | 0.512 | | 1.000 | | 0.999 |
| S3 |  |  | |  | |  | |  | |  | |  | |  | | 0.585 | | 0.721 |
| S4 |  |  | |  | |  | |  | |  | |  | |  | |  | | 0.999 |
| S5 |  |  | |  | |  | |  | |  | |  | |  | |  | |  |

**Table S2**: Statistical analysis of circadian timing between southern and northern *Nasonia vitripennis* under LD08:16. Indicated are *P*-values from ANOVA with a Tukey’s multiple-comparisons test. In bold *P* < 0.05.

|  | **Onset LD08:16** | | | | | | | | | | | | | | | | | | | | | | | | | | |
| --- | --- | --- | --- | --- | --- | --- | --- | --- | --- | --- | --- | --- | --- | --- | --- | --- | --- | --- | --- | --- | --- | --- | --- | --- | --- | --- | --- |
|  | N1 | | N2 | | | N3 | | | N4 | | N5 | | | S1 | | | S2 | | S3 | | | | S4 | | S5 | | |
| N1 |  | | 1.000 | | | 0.878 | | | 0.611 | | 0.376 | | | **< 0.001** | | | **0.014** | | **< 0.001** | | | | **0.003** | | **< 0.001** | | |
| N2 |  | |  | | | 0.891 | | | 0.60 | | 0.342 | | | **< 0.001** | | | **0.008** | | **< 0.001** | | | | **0.001** | | **< 0.001** | | |
| N3 |  | |  | | |  | | | 0.999 | | 0.999 | | | **0.030** | | | 0.507 | | **< 0.001** | | | | 0.202 | | **< 0.001** | | |
| N4 |  | |  | | |  | | |  | | 0.999 | | | 0.081 | | | 0.751 | | **< 0.001** | | | | 0.382 | | **< 0.001** | | |
| N5 |  | |  | | |  | | |  | |  | | | 0.157 | | | 0.899 | | **< 0.001** | | | | 0.569 | | **0.001** | | |
| S1 |  | |  | | |  | | |  | |  | | |  | | | 0.964 | | 0.765 | | | | 0.999 | | 0.897 | | |
| S2 |  | |  | | |  | | |  | |  | | |  | | |  | | 0.093 | | | | 0.999 | | 0.169 | | |
| S3 |  | |  | | |  | | |  | |  | | |  | | |  | |  | | | | 0.401 | | 0.999 | | |
| S4 |  | |  | | |  | | |  | |  | | |  | | |  | |  | | | |  | | 0.574 | | |
| S5 |  | |  | | |  | | |  | |  | | |  | | |  | |  | | | |  | |  | | |
|  | **Peak LD08:16** | | | | | | | | | | | | | | | | | | | | | | | | | | |
|  | N1 | N2 | | N3 | | | | N4 | | | | N5 | | | S1 | | | S2 | | | S3 | | | S4 | | | S5 |
| N1 |  | 1.000 | | **< 0.001** | | | | **< 0.001** | | | | **0.006** | | | **< 0.001** | | | **< 0.001** | | | **< 0.001** | | | **0.002** | | | **< 0.001** |
| N2 |  |  | | **< 0.001** | | | | **< 0.001** | | | | **< 0.001** | | | **< 0.001** | | | **0.009** | | | **< 0.001** | | | **0.001** | | | **< 0.001** |
| N3 |  |  | |  | | | | 1.000 | | | | 0.999 | | | 1.000 | | | 1.000 | | | **< 0.001** | | | 0.999 | | | **< 0.001** |
| N4 |  |  | |  | | | |  | | | | 0.997 | | | 0.999 | | | 1.000 | | | **< 0.001** | | | 0.999 | | | 0.999 |
| N5 |  |  | |  | | | |  | | | |  | | | 0.999 | | | 0.999 | | | **0.001** | | | 0.985 | | | **< 0.001** |
| S1 |  |  | |  | | | |  | | | |  | | |  | | | 1.000 | | | **< 0.001** | | | 0.999 | | | **< 0.001** |
| S2 |  |  | |  | | | |  | | | |  | | |  | | |  | | | **< 0.001** | | | 0.999 | | | **< 0.001** |
| S3 |  |  | |  | | | |  | | | |  | | |  | | |  | | |  | | | **< 0.001** | | | 0.709 |
| S4 |  |  | |  | | | |  | | | |  | | |  | | |  | | |  | | |  | | | **< 0.001** |
| S5 |  |  | |  | | | |  | | | |  | | |  | | |  | | |  | | |  | | |  |
|  | **Offset LD08:16** | | | | | | | | | | | | | | | | | | | | | | | | | | |
|  | N1 | N2 | | | N3 | | N4 | | | N5 | | | S1 | | | S2 | | | | S3 | | S4 | | | | S5 | |
| N1 |  | 0.999 | | | 0.156 | | 0.369 | | | **< 0.001** | | | **< 0.001** | | | **< 0.001** | | | | **< 0.001** | | **< 0.001** | | | | **< 0.001** | |
| N2 |  |  | | | **0.008** | | **0.04** | | | **< 0.001** | | | **< 0.001** | | | **< 0.001** | | | | **< 0.001** | | **< 0.001** | | | | **< 0.001** | |
| N3 |  |  | | |  | | 0.999 | | | **0.016** | | | **0.010** | | | **0.006** | | | | **< 0.001** | | 0.171 | | | | **< 0.001** | |
| N4 |  |  | | |  | |  | | | **0.006** | | | **0.004** | | | **0.002** | | | | **< 0.001** | | **0.079** | | | | **< 0.001** | |
| N5 |  |  | | |  | |  | | |  | | | 1.000 | | | 1.000 | | | | 0.316 | | 0.999 | | | | 0.697 | |
| S1 |  |  | | |  | |  | | |  | | |  | | | 1.000 | | | | 0.292 | | 0.998 | | | | 0.677 | |
| S2 |  |  | | |  | |  | | |  | | |  | | |  | | | | 0.413 | | 0.994 | | | | 0.799 | |
| S3 |  |  | | |  | |  | | |  | | |  | | |  | | | |  | | 0.053 | | | | 0.999 | |
| S4 |  |  | | |  | |  | | |  | | |  | | |  | | | |  | |  | | | | 0.211 | |
| S5 |  |  | | |  | |  | | |  | | |  | | |  | | | |  | |  | | | |  | |

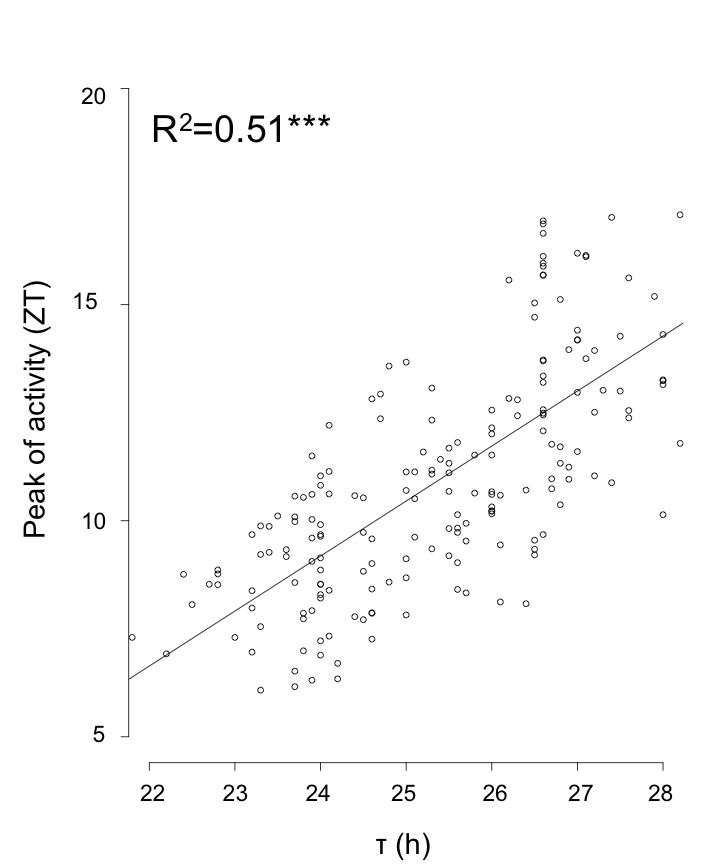

**Fig.S1 Correlation between peak of activity and free running period.** Free running period (τ) of southern and northern *Nasonia vitripennis*. Asterisks indicate a significant effect of activity timing on free running period (*** *P* < 2e-16, linear mixed effect model).
